# Genomic signatures of evolutionary rescue in bats surviving white-nose syndrome

**DOI:** 10.1101/470294

**Authors:** Sarah A. Gignoux-Wolfsohn, Malin L. Pinsky, Kathleen Kerwin, Carl Herzog, MacKenzie Hall, Alyssa B. Bennett, Nina H. Fefferman, Brooke Maslo

## Abstract

Rapid evolution of advantageous traits following abrupt environmental change can help populations grow and avoid extinction through evolutionary rescue. Here, we provide the first genetic evidence for rapid evolution in bat populations affected by white-nose syndrome (WNS). By comparing genetic samples from before and after little brown bat populations were decimated by WNS, we identified signatures of soft selection on standing genetic variation. This selection occurred at multiple loci in genes linked to hibernation behavior rather than immune function, suggesting that differences in hibernation strategy have allowed these bats to survive infection with WNS. Through these findings, we suggest that evolutionary rescue can be a conservation-relevant process even in slowly reproducing taxa threatened with extinction.

---

Many North American bat species are currently facing extinction due to white-nose syndrome (WNS), an infectious disease caused by the introduced fungal pathogen *Pseudogymnoascus destructans (1, 2). P. destructans* infects bats during hibernation; it creates lesions in wing membranes, disrupts homeostasis and depletes energy stores by altering metabolism and increasing the frequency of arousal from torpor *(1, 3)*. Some remnant populations of one species, the little brown bat *(Myotis lucifugus)*, have returned to neutral or positive growth in recent years after abundance declines of up to 98% *(4)*. Furthermore, some populations are beginning to display the classic U-shaped trajectory associated with evolutionary rescue *(5, 6)*. Recent analyses of population declines and infection rates suggest that host resistance is the driving mechanism behind these population trajectories, rather than alternative mechanisms such as host tolerance, demographic compensation, or reduced pathogen virulence *(7)*. What remains unclear is whether this resistance results from the acquisition of adaptive immunity or from selection of genetically resistant individuals *(6)*. Only the latter mechanism is consistent with evolutionary rescue.

To identify WNS-induced genetic changes, we performed whole genome sequencing on 132 individuals from four hibernating colonies (hibernacula) within the epicenter of WNS emergence (Northeastern US, Fig. 1A). Two hibernacula, Hibernia Mine (Hibernia) and Walter Williams Preserve (Williams), were sampled both pre-infection by WNS (2008 and 2009, respectively) and post-infection by WNS (2016, Fig. 1B). One hibernaculum, Greeley Mine (Greeley), was sampled pre-WNS (2008); and another, Aeolus Cave (Aeolus), was sampled post-WNS (2016). We used probabilistic methods that explicitly account for depth of coverage (Table S1) to identify a total of 31,517,948 SNPs (single nucleotide polymorphisms) and to calculate allele frequencies through time for SNPs across the genome.

**Fig. 1.**
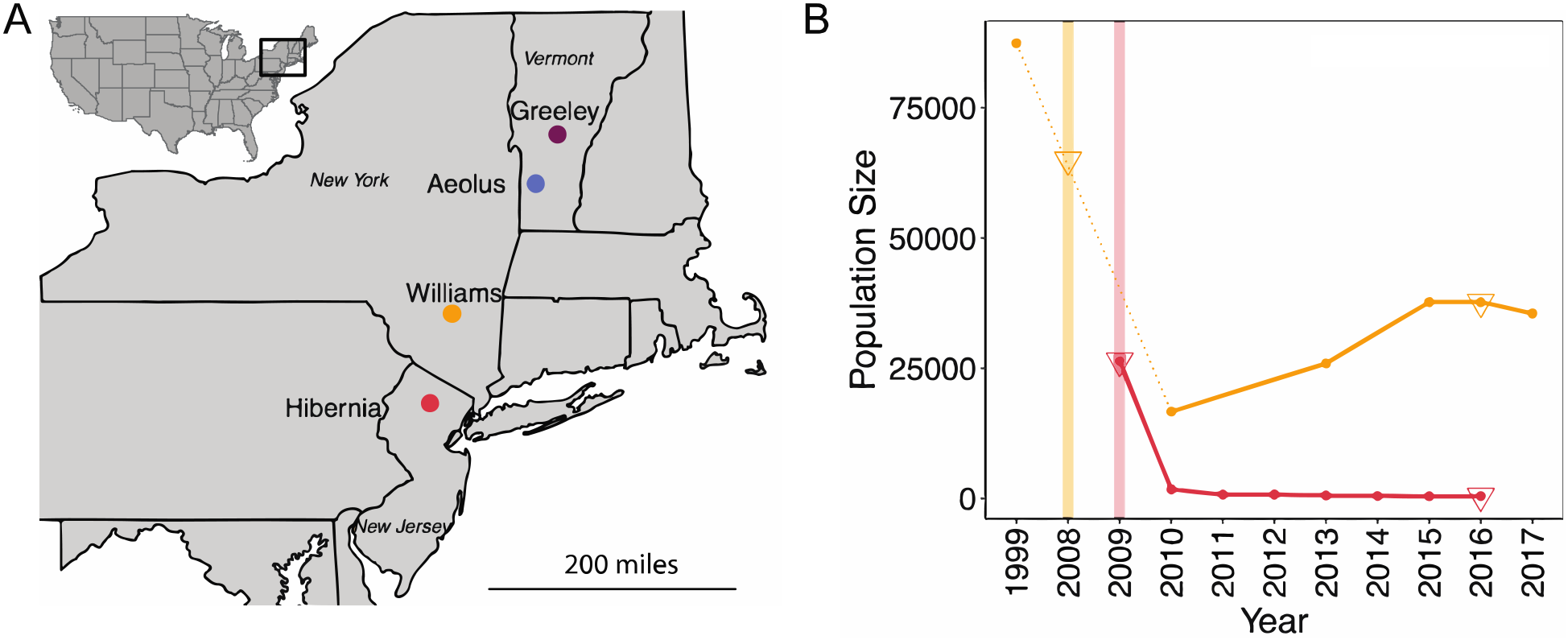
Four hibernacula were sampled across time and are experiencing different rates of population recovery. A) Location of hibernacula sampled in the northeast United States. B) Population trajectories for Williams and Hibernia, color coded according to A). Vertical lines represent the time of initial reports of WNS for each hibernaculum. The dotted line represents missing data for Williams because population estimates were not made between 1999 and 2010 (1999 is the presumed pre-WNS baseline). The triangles indicate sampling points for each hibernaculum. Population estimates were not available for Greeley and Aeolus.

We found little geographic structure at a genome-wide level, including low pairwise F_ST_ values (pre-WNS samples 0.0176 +/− 0.000033, post-WNS samples 0.0143 +/− 0.000333) and weak population structure that did not correspond to geography (Fig. 2A, B). A lack of population structure is consistent with previous studies demonstrating near-panmixia over evolutionary time in bat populations east of the Rocky Mountains *(8, 9)*, likely because even a small number of bats moving between hibernacula reduces population structure on an evolutionary timescale *(10).* Despite near-panmixia over evolutionary time, most bats have fidelity to mating sites and hibernacula on ecological timescales (11). Admixture proportions indicated that two of the bats from Williams may have originated from a second evolutionary population (Fig. 2B). To avoid confounding population changes with signatures of selection, we removed these two bats, plus two other potentially genetically divergent bats (Fig. 2A), from calculations of allele frequency for the results presented in the main paper (Fig. 4, but see Fig. S4 for results with all bats included). Further, we found little genome-wide temporal structure as a result of WNS (F_ST_ comparing pre-WNS populations to post-WNS populations, Hibernia: 0.016, Williams: 0.017). We therefore do not find any genetic evidence to suggest that pre-WNS populations were replaced by individuals from other populations (Fig. 2).

**Fig. 2.**
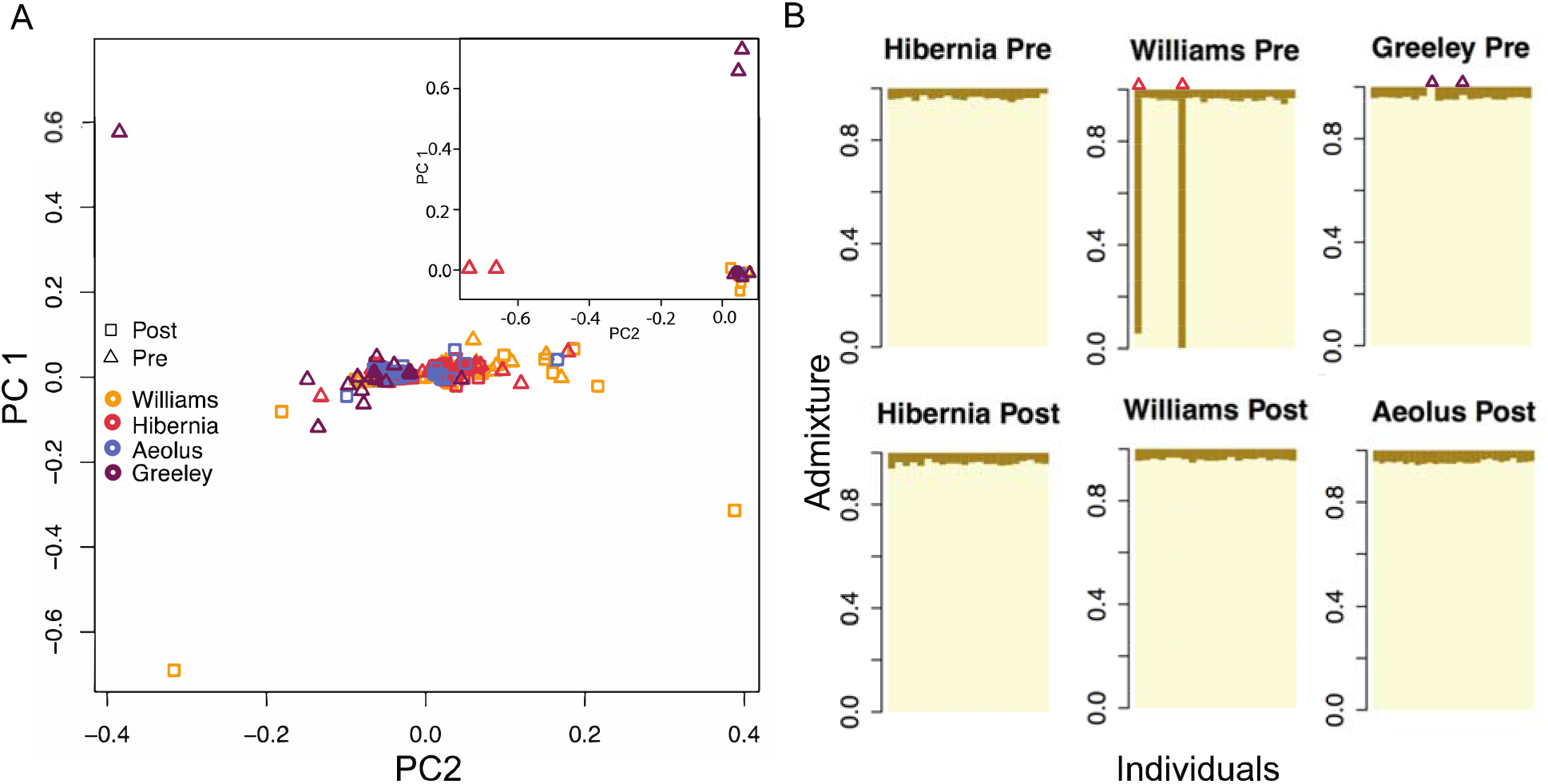
Population structure of little brown bats in the northeastern United States does not correspond to geography or time. A) PCA plot shows four potential genetically divergent bats (inset) and a lack of population structure in the majority of individuals (with divergent bats removed, main figure). MAP analysis identified one significant principal component (PC 1). B) Admixture plot showing two populations, which was the most likely value determined by MAP analysis. Colors denote different inferred ancestral origin populations. The four potentially divergent bats from the PCA are marked at the top of the admixture plots.

Demographic declines were dramatic within the smaller Hibernia colony, with an estimated population size of 26,438 individuals in 2009, declining by 98% from 2009-2015, and returning to pre-WNS survival rates and a slow but positive population growth rate after 2015 (Fig. 1B, Fig. S1). Steep but less dramatic declines from 87,401 individuals in 1999 to 16,673 individuals in 2010 (80% decline) were followed by partial recovery of the population at the larger Williams colony (Fig. 1B). Tajima’s D is a genetic statistic that typically increases after a population bottleneck as rare variants are lost (and intermediate frequency alleles become more common). The effects tend to be strongest in populations that reach the smallest sizes *(12).* We observed that Tajima’s D increased through time in the smaller and more heavily bottlenecked Hibernia population, but decreased slightly in the relatively more abundant and rapidly recovering Williams population (Mann-Whitney U test p<0.001 for both populations, Fig. 3).

**Fig. 3.**
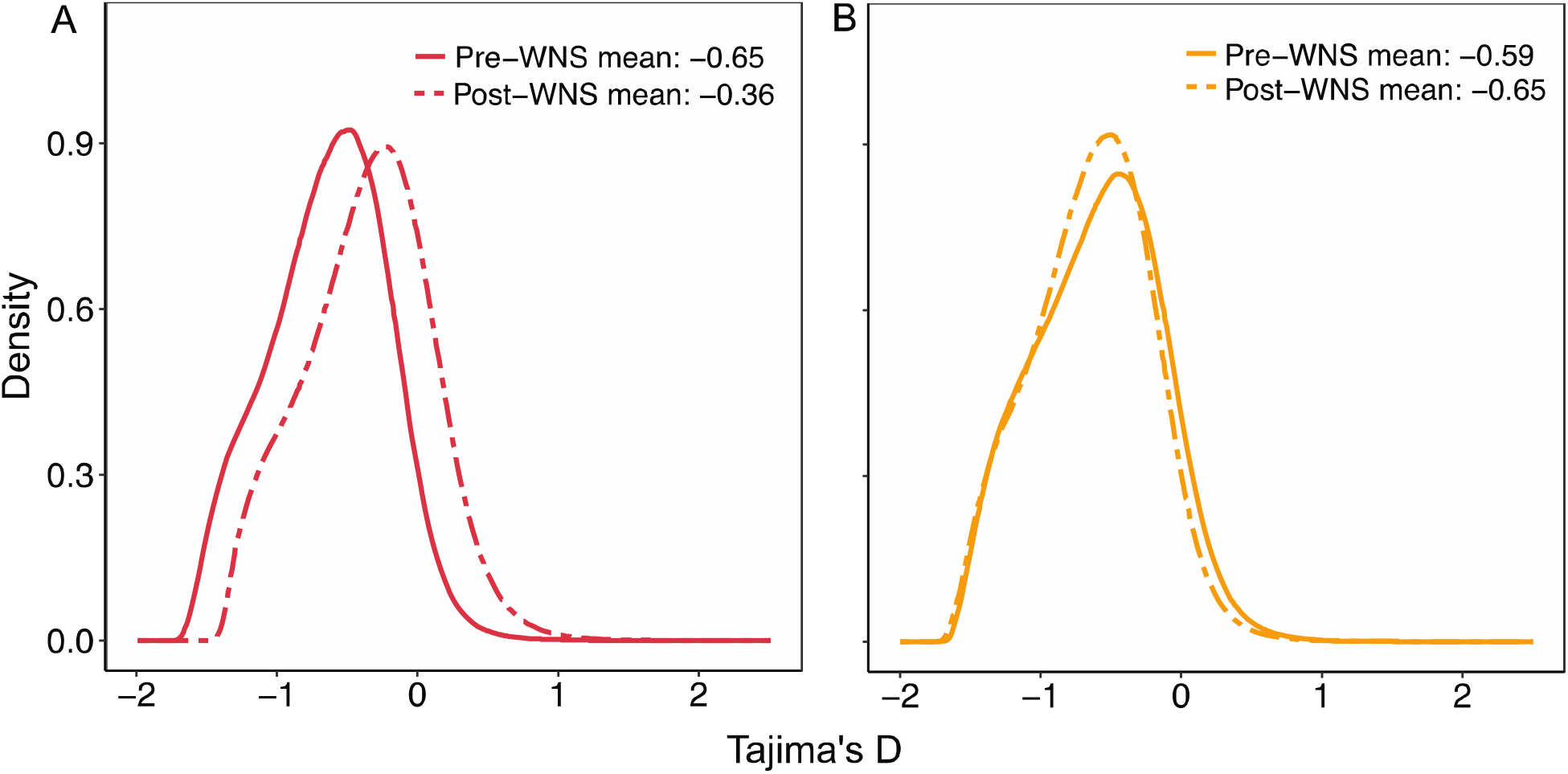
Change in Tajima’s D reflects differences in population declines in Hibernia and Williams. Density plots of Tajima’s D calculated in sliding windows across the genome in A) Hibernia and B) Williams from samples pre-(solid line) and post-(dotted line) infection with WNS. Mean Tajima’s D across the entire genome is noted for each hibernaculum.

To test for parallel signals of selection in populations from Hibernia and Williams, we compared allele frequency changes through time against a null model of genetic drift and sampling variance. This approach identified 62 SNPs with significantly greater allele frequency changes than would be expected from genetic drift and sampling variance alone (Table S5, Fig. 4A, B). The candidate SNPs were present at low to moderate frequencies in pre-WNS populations from both Hibernia and Williams and at low to intermediate frequencies in the pre-WNS Greeley population (Fig. 4D). These starting allele frequencies are consistent with a soft selective sweep from standing genetic variation, a likely method of selection given the short timescale of WNS infection, mortality, and recovery. The short timeframe makes novel mutation-derived hard sweeps unlikely. Soft selection has recently been proposed as a common and possibly dominant method of selection, though one that has been largely overlooked by genome scans because detection is difficult when temporal genetic data are lacking *(13).* Sampling the same sites through time therefore provides a method for detecting selection that would otherwise be nearly invisible *(14)*.

**Fig. 4.**
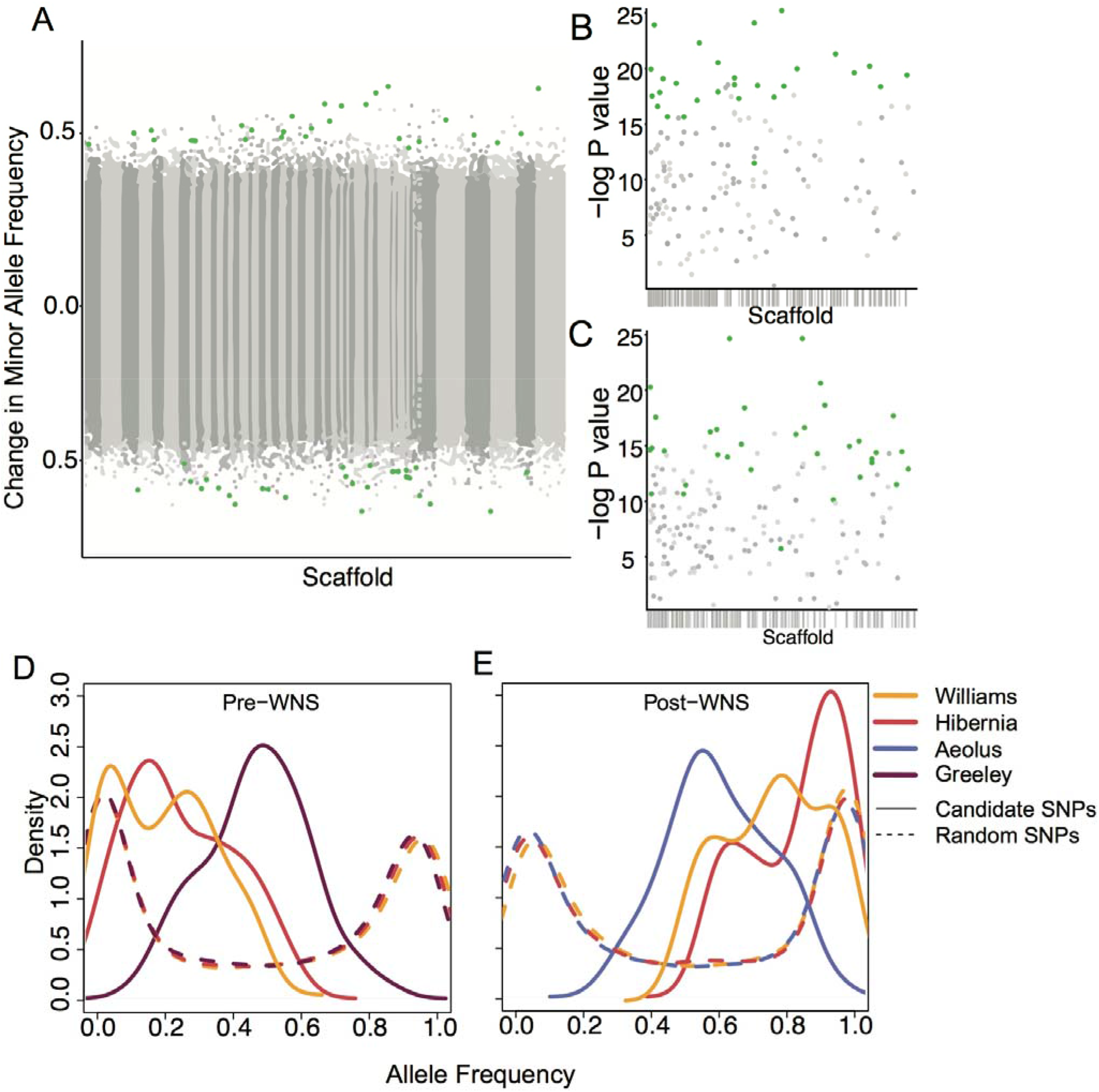
SNPs with signals of soft selection were located throughout the genome, were present at low to intermediate frequencies prior to WNS infection, and swept to incomplete or nearly complete fixation after WNS. A) Change in allele frequency between pre- and post-WNS samples from Hibernia and Williams. Scaffolds are displayed in the order of the MyoLuc 2.0 assembly. Significant SNPs at FDR=0.2 are highlighted in green. B, C) −log10 P-values for SNPs with a positive (B) or negative (C) change in allele frequency greater than 0.5 in both Hibernia and Williams. Scaffolds are colored along x axis. Significant SNPS are highlighted in green. D) Density plots of allele frequencies for significant candidate SNPs (solid lines) and for SNPs randomly chosen throughout the genome (dotted lines) at each site pre-WNS and E) post-WNS.

Post-WNS allele frequencies in Hibernia, Williams and Aeolus ranged from intermediate to high, suggesting incomplete selective sweeps at some loci and near-complete sweeps at others (Fig. 4E). Pre-WNS frequencies of candidate SNPs were highest in Williams and post-WNS frequencies were lowest at Aeolus, which suggests that—while some degree of parallel selection is occurring across multiple populations—selection is not acting to the same degree on the same loci in all populations affected by WNS. Such signatures are consistent with soft selection for traits controlled by many loci *(13).* As further evidence against hard selection, genomic regions surrounding SNPs putatively under selection did not show classic signatures of hard selection, such as changes in π and Tajima’s D (Figs. S2 and S3).

We further tested whether parallel changes in allele frequencies across Hibernia and Williams could be explained by dispersal between populations. However, we found that migrating individuals would have to make up >52% of each population (Table S4). Mark-recapture data collected at Hibernia over the last seven years found that 95% of bats banded at this hibernaculum were subsequently recaptured in later years (Table S3). Furthermore, biannual surveys of Williams have shown that <0.02% of bats at Williams (out of 16,000) have been detected migrating from Hibernia. No bats from Williams have been found at Hibernia (Tables S2 and S4). This result suggests that selection acted *in situ* and largely independently in each population after WNS infection.

Sixteen of the 62 candidate SNPs under selection were located in annotated regions of the genome (Table S5). These SNPs were all located in presumed introns. Introns can play important regulatory roles, and mutations in introns have been associated with numerous diseases *(15).* Introns have a higher mutation rate due to lower evolutionary constraint and may be good candidates for selection on standing genetic variation.

Functional annotation suggested that selection has acted on hibernation behavior, with most (10 of 16) annotated genes involved in regulating processes associated with hibernation (Table S6). Three genes (SORCS3, PCDH17, and KREMEN1) were associated with neural regeneration and plasticity and may regulate the neural changes involved in hibernating mammals: shrinking of neurons, inhibition of neurogenesis, and reduction of synapses, all of which are rapidly reversed upon arousal from hibernation *(16).* Hibernating mammals have been proposed as a model system for both neuroprotection *(17)* and reversing neurodegenerative diseases *(18).* Genome wide association studies have linked many of these genes to human neurodegenerative diseases such as Parkinson’s, Alzheimer’s, and neuratypical phenotypes (Table S6), though their particular function in bats remains unclear.

We identified several SNPs in genes associated with insulin regulation and fat accumulation (REPS2, SOX5, ADCY3, CMIP; Table S6). Hibernating bats undergo significant changes in insulin levels, fat accumulation, and metabolism throughout the hibernation cycle *(19, 20)*, essentially experiencing a reversible form of type 2 diabetes *(21)*. Although slightly below our significance threshold, it is important to note that we identified a SNP with allele frequency changes greater than 0.45 in both Hibernia and Williams populations, low frequency in (pre-WNS) Greely and high frequency in (post-WNS) Aeolus within a gene previously associated with ground squirrel hibernation (EXOC4) *(22).* Contrary to other studies of disease-induced selection, we found only one candidate SNP in a gene involved in immune system function (MASP1, part of the complement pathway).

When hibernating bats are infected with *P. destructans*, the frequency of torpor arousal increases and metabolism is altered (23), which prematurely depletes fat stores and leads to mortality *(3, 23).* Selection on hibernation-related neural and metabolic genes in this host-pathogen system is therefore a logical but unintuitive consequence of WNS infection. Disease-induced arousal may be a failed attempt to fight off the pathogen by re-activating the immune system *(24)*, but an increased immune response and production of anti-*P. destructans* antibodies is not correlated with survival and is therefore likely only reducing energy reserves *(25).* Furthermore, European bats that harbor *P. destructans* but do not display signs of WNS frequently have lower levels of anti-*P. destructans* antibodies than uninfected bats (26). Intriguingly, big brown bats *(Eptesicus fuscus)*, which have suffered much smaller population declines, increase their torpor duration when exposed to *P. destructans (27).* Our results suggest that differences in hibernation strategy may be the key to host survival and that evolutionary rescue may have altered hibernation behavior in little brown bats. Two of the genes identified here (PCDH17 and REPS2) have been previously found to be differentially expressed in bat wing tissue when infected with *P. destructans (28).* Further research will be needed to directly link these genetic changes to changes in gene expression and hibernation phenotypes.

Surviving individuals may have the potential to rescue little brown bat populations and ultimately return them to stable population levels *(6)*. Characterizing the full suite of adaptive alleles present will require detecting SNPs likely under selection in other populations given that differences in pre-WNS genetic variation determines which alleles are available for selection. Management decisions *(e.g.*, whether to deploy treatments for *P. destructans* or to protect populations from other detrimental factors) can be informed by knowing the frequency of resistant individuals in both uninfected and infected populations. The deployment of vaccines or treatments for *P. destructans* may be most needed in populations with low evolutionary potential *(29).* This study suggests that evolutionary rescue may be occurring in other instances of pathogen-induced selection. By closely observing wild populations of non-model organisms, we can document unique natural experiments that help to expand our understanding of how and when natural selection occurs.

## Materials and Methods

### Mark-recapture

We began mark-recapture research at Hibernia Mine in 2010, one year after white-nose syndrome (WNS) emergence at the site. During annual surveys from 2010-2017, we captured and marked 1,262 (476 females, 786 males) little brown bats with unique 2.9-mm or 2.4-mm lipped alloy bands (Porzana Ltd., Icklesham, UK). We classified all bats as adults, as it is not possible to accurately determine age class after an individual’s first summer.

Population monitoring activities were carried out under the authority of a cooperative agreement between the New Jersey Division of Fish and Wildlife (formerly NJ Division of Fish, Game and Shellfisheries) and the US Fish and Wildlife Service dated June 23, 1976. National White-nose Syndrome Decontamination Protocols *(30)* were followed during all visits. We analyzed encounter histories of individuals using Cormack-Jolly-Seber (CJS) models in Program MARK (31) and examined the effects of sex, year, and time since WNS arrival on annual survival. We developed 24 *a priori* candidate models containing combinations of constant, yearly, time trend, and sex-specific effects on annual survival and recapture probabilities, including a global model that included time-dependent and sex-specific survival and recapture. To test for goodness of fit, we used a parametric bootstrapping procedure with 500 simulations of our penultimate model and calculated a variance inflation factor of *ĉ* = 1.26. Because we detected slight overdispersion in our data, we used small-sample corrected Quasi-Akaike’s Information Criterion (QAIC_c_) adjusted by *ĉ* = 1.26 (32). We ranked candidate models by ΔQAIC_c_ and Quasi-Akaike weights (*w*), which represent the relative likelihood of the model, given the data *(33)*. To reduce model selection bias and uncertainty, we averaged all models contributing to cumulative *w* ≥ 0.85 (top five models) and calculated parameter estimates based on weighted averages of the parameter estimates in the top models *(32, 34)*. Model results confirmed previously published findings *(35)* of a linear increase in annual survival with time since WNS arrival. Recapture rates averaged ~0.40 for both sexes.

To determine the post-WNS growth rate of the remnant Hibernia population, we incorporated our survival estimates into seven 2-stage Lefkovitch matrices *(35–37)*. Each matrix represented one study year from 2010-2016:

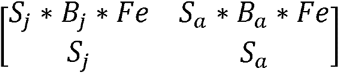

where *S* represents survival of female little brown bats; *B* represents the probability that a female returns to a maternity colony to breed; and subscripts *j* and *a* indicate values for juveniles or adults, respectively. Little brown bats have a single pup each year and are sexually mature by the end of their first summer, so we held fecundity (Fe) constant for both age classes at *Fe* = 1. We assigned probability of breeding for adults and juveniles to values of *B_j_* = 0.38 and *B_a_* = 0.85, respectively, based upon published estimates generated from either 15 years of pre-WNS mark-recapture data or post-WNS adult reproductive rates *(2, 38, 39)*. We fixed juvenile survival as a constant proportion of adult survival (Sj = 0.47 * Sa; *2*). We then calculated the dominant eigenvalue of each matrix. We projected the Hibernia little brown bat population through the 2010-2016 matrices and then continued the projection an additional 93 years using the 2016 matrix. We ran a 10,000-iteration Monte Carlo simulation, allowing *S_a_* and *B_j_* to vary stochastically based on random number generation from beta distributions specified from the means and variances of each survival parameter. From this procedure, we generated a mean stochastic yearly growth rate of λ = 1.03.

### Sample collection

We collected tissue samples at two timepoints, before (pre-WNS) and after (post-WNS) disease-induced population declines. We collected pre-WNS samples from frozen specimens collected in the first year of disease emergence at Walter Williams Preserve, Ulster County, New York (2008, n=23); Hibernia Mine, Morris County, New Jersey (2009, n=19); and Greeley Mine, Windsor County, Vermont (2009, n=19). We collected post-WNS samples from live individuals within the remnant populations at Hibernia Mine (n=22) and Walter Williams Preserve (n=20), and from Aeolus Cave (Bennington County, Vermont, n=26) in 2016. At the time of post-WNS sampling, Greeley Mine was inaccessible. Sample collection followed Simmons (2008). Briefly, a 2-mm wing tissue biopsy was taken (Miltex) and placed in RNALater (Ambion). Samples were collected under IACUC Protocol #15-068 (Rutgers University) and appropriate state permits.

### Library preparation and sequencing

DNA was extracted using the QIAGEN Blood and Tissue kit with the addition of RNase A to remove RNA contamination and following recommendations to increase the concentration of DNA (10 min elution step, smaller volume of elution buffer). Fragmented DNA was identified using 1.5% gel electrophoresis and when fragments <1 kbp were present, a cleanup step was performed with AMPure XP beads (Agencourt). DNA concentration was measured using a Qubit HS DNA Assay (Invitrogen).

Library prep was performed using the Illumina Nextera kit following the low-coverage whole genome sequencing protocol developed by Therkildsen and Palumbi (2016), modified from Baym *et al.* (2015). Briefly, samples were fragmented and adapters were added using tagmentase in a 2.5 μl volume. Because the ideal concentration of DNA given the size of the *M. lucifugus* genome was determined to be 10 ng/μl, which was above the concentration for many of our samples, we added an additional step to dilute the tagmentase for low concentration samples using 10xTB.

A two-step PCR procedure was then conducted using the KAPA Library Amplification kit and the Nextera index kit (primers N517-N504 and N701-N706) with a total of 12 cycles (8+4) to add a unique combination of barcodes to each sample and to amplify the resulting libraries. We then purified and size selected these products with AMPure XP beads, quantified the concentration of each library using a Qubit HS DNA Assay, and examined the fragment size using an Agilent BioAnalyzer chip. Finally, we pooled the resulting libraries together in equal concentrations by mass. These pools were sequenced in 7 rapid runs (each of 2 lanes/run) on the Illumina HiSeq 2500 at the Princeton University Lewis-Siegler Institute. We achieved an average depth of coverage of 2.42 +/− 1.24. This depth of coverage allowed us to call SNPs and calculate population-level allele frequencies using established methods that take genotype uncertainty into account *(43, 44)*. We did not need nor attempt to call individual genotypes. Individual-based but low coverage approaches such as these have increased efficiency, precision, and cost-effectiveness for estimating allele frequencies as compared to higher coverage approaches *(41, 45)*, which can place undue confidence in what are actually sequencing or PCR errors. Furthermore, this approach eliminates non-equimolar DNA errors or stochastic amplification differences associated with Pool-seq approaches *(46, 41, 47)*.

### Code Availability

All scripts and notebooks can be found at https://github.com/sagw/WNS_WGS/ and relative paths referenced below are to be appended to this base path.

### Bioinformatics

To remove exact duplicate sequences (likely optical duplicates), we ran FastUniq in paired-end mode. We then used Trimmomatic v0.36 in paired-end mode to remove Illumina adapters, remove reads with an average phred score below 33, trim any reads where a 4 basepair sliding window phred score fell below 15, and discard trimmed reads shorter than 30 basepairs.

We filtered contaminant sequences using fastqscreen v0.95 (https://www.bioinformatics.babraham.ac.uk/projects/fastq_screen/) with Bowtie2. Reads were screened against the human genome (ch38), all bacterial genomes, all fungal genomes including *P. destructans*, and all viral genomes available from NCBI (ftp://ftp.ncbi.nlm.nih.gov/genomes/genbank/). All reads were tagged for contaminants using fastqscreen. Less than 10% of reads were removed from unpaired reads using fastqscreen in filter mode and from paired reads using a custom script (Scripts/pfilter.py).

We mapped resulting paired and unpaired reads to the *Myotis lucifugus* 2.0 genome (ftp://ftp.ensembl.org/pub/release-87/fasta/myotis_lucifugus/dna/) using Bowtie2 in very sensitive local mode (48). We then sorted sam files, converted to bam, and removed duplicate sequences using Picard (https://broadinstitute.github.io/picard/) (MarkDuplicates). We used Bedtools *(49)* to calculate percent coverage and discarded samples with reads that covered <20% of the genome. These filters resulted in 132 samples with an average coverage of 0.5x. we used GEM *(50)* to determine mappability of the reference genome. Only regions with a mappability score of 1 were used (1.7 out of 2.2 gigabases). Finally, we removed repeat regions as determined using RepeatMasker *(51)*.

### SNP calling and summary statistics

Following previous studies using low-coverage data (e.g., *(52), (53)*), we called SNPs over all samples using ANGSD (v0.92) with the following parameters: the samtools model was used to estimate genotype likelihoods from the mapped reads (-GL 1), major and minor alleles and frequency were estimated from genotype likelihoods (-doMaf 1 -doMajorMinor 1). We used the following quality filters (requires -doCounts 1): a minimum quality score of 20, minimum mapping score of 30, minimum number of individuals with data of 68 and max depth over all individuals of 1320. We used a cut-off *p*-value of 10^-6^ (Likelihood Ratio Test). See Notebooks/Angsd_all_SNPs for all details.

We calculated genotype likelihoods for called SNPs with ANGSD in beagle format (-doGlf 2). These likelihoods were used with PCAngsd to 1) calculate a covariance matrix 2) determine the number of significant principal components (D) using Shriner’s implementation of Velicer’s minimum average partial (MAP) test, and 3) determine admixture proportions using nonnegative matrix factorization with the number of ancestral populations (K) calculated as D+1 *(54).* The PCA and admixture proportions were then visualized using R (see Notebooks/Angsd_all_SNPs and Notebooks/Angsd_all_PCA_graphs).

In order to calculate FST between subpopulations, we first calculated unfolded sample allele frequencies with ANGSD using the reference genome to polarize the alleles for each subpopulation (-doSaf 1). We then created the following joint site frequency spectra with the ANGSD program realSFS: Hibernia pre-post, Williams pre-post, Hibernia-Williams-Greeley pre, Hibernia-Williams-Aeolus post. An average FST was calculated across the entire genome and in a sliding window across the genome. See Notebooks/Angsd_all_FST.

We used the per-population sample allele frequencies to create unfolded site frequency spectra for each population. The sample allele frequencies and site frequency spectra were then used to calculate θ (population mutation rate) across the genome (-doThetas 1), and then calculate θ_w_, π, and Tajima’s D both across the genome and in a sliding window with the ANGSD program thetaStat (do_stat -win 1000 -step 1 -type 1). See Notebooks/Angsd_all_thetas.

We estimated minor allele frequencies for each called SNP in each subpopulation using the previously inferred major and minor alleles (-doMajorMinor 3). Allele frequency was summed over 3 possible minor alleles weighted by probabilities (-doMaf 2). Maximum depth was set at 270. See Notebooks/Angsd_all_SNPs.

### Identifying SNPs putatively under selection

For the Hibernia and Williams colonies (excluding outlier bats, see Fig. 1), change in minor allele frequency was calculated from MAFs (method 2) as maf_pos_t-maf_pre_. We considered SNPs with a high change in allele frequency as those with changes greater than 0.5 or less than −0.5 in both Hibernia and Williams (in the same direction). To separate SNPs with high allele frequency changes due to sampling error or genetic drift from those under selection, we developed locus-specific null models for allele frequency change. We first accounted for sampling error by using ANGSD to calculate minor allele frequencies from 100 bootstrapped file lists for Hibernia pre-WNS and post-WNS and Williams pre-WNS and post-WNS. Wright-Fisher simulations (1000 for each value) were then performed to estimate genetic drift over the two generations that have elapsed between pre- and post-WNS samples *(55, 56)*. We assumed the two generations had effective population sizes of 424 and 296, respectively. These values were calculated from the more reliable Hibernia census data using method described by Waples *(57)*, where average lifespan is 15 years, age to maturity is 1 year and generation time is 6 years *(38)*. Hibernia has a smaller population size and using these data likely overestimated drift and made our test more conservative. *P*-values were calculated for each SNP as *p*=(*r*+1)/(*n*+1), where *r* was the number of times the simulated post-WNS allele frequencies were greater than sampled bootstrapped post-WNS allele frequencies, and *n* was the total number of simulations *(58)*. All analyses can be found in Notebooks/All_AlleleFreqChange.ipynb.

Because there is no evidence of significant bat migration between colonies in the time frame of this study (even if populations were panmictic in evolutionary time), the NJ and NY populations were considered independent and combined *p*-values were calculated using the sum log method in the R package metap (59). To account for multiple hypothesis testing, significance was then determined using the Benjamini-Hochberg method. Values were ranked from smallest to largest and the largest *p*-value with the *p*<(i/m)Q was used as the significance cut off, where i was rank, m was the number of SNPs in both Hibernia and Williams, and Q was our chosen false discovery rate of 0.2.

The immigration rate that would be required to achieve the observed allele frequency changes was calculated as % immigrants = 100*(Fpost1-Fpre1)/(Fpost2-Fpre1), where % immigrants is percent immigrants in population 1 post-WNS, Fpost1 was allele frequency in population 1 post-WNS, Fpre1 was allele frequency in population 1 pre-WNS, Fpost2 was allele frequency in population 2 post-WNS.

The NCBI *Myotis Lucifugus* Annotation Release 102 was used to identify putative genes that contained the candidate SNPs and the Ensembl Variant Effect predictor was used to identify the location of SNPs within genes.

## Acknowledgements

The authors would like to thank Nina Therkildsen, Noah Rose, Michelle Stuart, and Wei Wang for help with troubleshooting; Chris Gignoux, Lisa Komoroske, Jennifer Hoey, Katrina Catalano, Winifred Frick, A. Marm Kilpatrick, and Erik Holum for discussions about bat ecology and data analyses; Chris Gignoux, Maarten Vonhof, and Katrina Lohan for reviewing a draft manuscript. We thank Brian Schumm and multiple other field personnel for assistance in sampling bats.

## Author contributions

BM, MP, and NF conceived of the idea, BM, KK, CH, MH, AB performed sample collection, SGW conducted all lab work and data analysis with input from BM and MP, SGW wrote the manuscript with input from all coauthors.

## Data and materials availability

Sequences are available from the short read archive (bioproject: PRJNA509256, accession #s SAMN10574961-SAMN10575092). All scripts and notebooks can be found at https://github.com/sagw/WNS_WGS/

## References and Notes

1. A. Gargas, M. T. Trest, M. Christensen, T. J. Volk, D. S. Blehert, Geomyces destructans sp. nov. associated with bat white-nose syndrome. Mycotaxon 108, 147–154 (2009).

2. W. F. Frick et al., An emerging disease causes regional population collapse of a common North American bat species. Science 329, 679–682 (2010).

3. D. M. Reeder et al., Frequent arousal from hibernation linked to severity of infection and mortality in bats with white-nose syndrome. PLoS One 7, e38920 (2012).

4. W. Frick et al., Pathogen dynamics during invasion and establishment of white-nose syndrome explain mchanisms of host persistence. Ecology 98, 624–631 (2017).

5. E. Vander Wal, D. Garant, M. Festa-Bianchet, F. Pelletier, Evolutionary rescue in vertebrates: evidence, applications and uncertainty. Philosophical transactions of the Royal Society of London. Series B, Biological sciences 368, 20120090 (2013).

6. B. Maslo, N. H. Fefferman, A case study of bats and white-nose syndrome demonstrating how to model population viability with evolutionary effects. Conservation biology: the journal of the Society for Conservation Biology 29, 1176–1185 (2015).

7. K. E. Langwig et al., Resistance in persisting bat populations after white-nose syndrome invasion. Philosophical transactions of the Royal Society of London. Series B, Biological sciences 372,(2017).

8. A. P. Wilder, T. H. Kunz, M. D. Sorenson, Population genetic structure of a common host predicts the spread of white-nose syndrome, an emerging infectious disease in bats. Molecular ecology 24, 5495–5506 (2015).

9. M. J. Vonhof, A. L. Russell, C. M. Miller-Butterworth, Range-Wide Genetic Analysis of Little Brown Bat (Myotis lucifugus) Populations: Estimating the Risk of Spread of White-Nose Syndrome. PLoS One 10, e0128713 (2015).

10. R. S. Waples, O. Gaggiotti, What is a population? An empirical evaluation of some genetic methods for identifying the number of gene pools and their degree of connectivity. Molecular ecology 15, 1419–1439 (2006).

11. K. J. O. Norquay, F. Martinez-Nuñez, J. E. Dubois, K. M. Monson, C. K. R. Willis, Long-distance movements of little brown bats *(Myotis lucifugus)*. Journal of Mammalogy 94, 506–515 (2013).

12. Tajima, The Effect of Change in Population Size on DNA Polymorphism. Genetics 123, 597–601 (1989).

13. P. W. Messer, D. A. Petrov, Population genomics of rapid adaptation by soft selective sweeps. Trends Ecol Evol 28, 659–669 (2013).

14. M. Przeworski, G. Coop, J. D. Wall, The Signature of Positive Selection on Standing Genetic Variation. Evolution; international journal of organic evolution 59, 2312 (2005).

15. M. Ma et al., Disease-associated variants in different categories of disease located in distinct regulatory elements. BMC genomics 16,(2015).

16. G. Leon-Espinosa et al., Decreased adult neurogenesis in hibernating Syrian hamster. Neuroscience 333, 181–192 (2016).

17. F. Zhou et al., Hibernation, a Model of Neuroprotection. American Journal of Pathology 158,(2001).

18. N. Braidy et al., Recent rodent models for Alzheimer’s disease: clinical implications and basic research. J Neural Transm (Vienna) 119, 173–195 (2012).

19. H. V. Carey, M. T. Andrews, S. L. Martin, Mammalian Hibernation: Cellular and molecular Responses to Depressed Metabolism and Low Temperature. Physiol Rev 83, 1153–1181 (2003).

20. W. A. Bauman, Seasonal Changes in Pancreatic Insulin and Glucagon in the Little Brown Bat Pancreas 5, 342–346 (1990).

21. C. W. Wu, K. K. Biggar, K. B. Storey, Biochemical adaptations of mammalian hibernation: exploring squirrels as a perspective model for naturally induced reversible insulin resistance. Brazilian Journal of Medical and Biological Research 46, 1–13 (2013).

22. K. R. Grabek et al., Genetic architecture drives seasonal onset of hibernation in the 13-lined ground squirrel. (2017).

23. M. L. Verant et al., White-nose syndrome initiates a cascade of physiological disturbances in the hibernating bat host. BMC Physiology 14,(2014).

24. K. B. Storey, J. M. Storey, in Functional Metabolism: Regulation and Adaptation, K. B. Storey, Ed. (John Wiley & Sons, 2004).

25. T. M. Lilley et al., Immune responses in hibernating little brown myotis (Myotis lucifugus) with white-nose syndrome. Proceedings of the Royal Society B: Biological Sciences 284, 20162232 (2017).

26. J. S. Johnson et al., Antibodies toPseudogymnoascus destructansare not sufficient for protection against white-nose syndrome. Ecology and evolution 5, 2203–2214 (2015).

27. M. S. Moore et al., Energy Conserving Thermoregulatory Patterns and Lower Disease Severity in a Bat Resistant to the Impacts of White-Nose Syndrome. J Comp Physiol B 188, 163–176 (2018).

28. K. A. Field et al., Effect of torpor on host transcriptomic responses to a fungal pathogen in hibernating bats. Molecular ecology, (2018).

29. B. Maslo, S. Gignoux-Wolfsohn, N. H. Fefferman, Success of Wildlife Disease Treatment Depends on Host Immune Response. frontiers in ecology and evolution, (2017).

31. G. C. White, K. P. Burnham, Program MARK: Survival Estimation from populations of marked animals. Bird Study 46,(1999).

32. K. P. Burnham, D. R. Anderson, Model selection and multimodel inference: a practical information-theoretic approach., (Springer, 2002).

33. J. B. Johnson, K. S. Omland, Model selection in ecology and evolution. Trends in ecology & evolution 19, 101–108 (2004).

34. K. P. Burnham, D. R. Anderson, K. P. Huyvaert, IC model selection and multimodel inference in behavioral ecology: some background, observations, and comparisons. Behavioral Ecology and Sociobiology 65, 23–35 (2011).

35. B. Maslo, M. Valent, J. F. Gumbs, W. Frick, Conservation implications of ameliorating survival of little brown bats with white-nose syndrome. Ecological Applications 25, 1832–1840 (2015).

36. H. Caswell, Matrix population models., (Wiley Online Library, 2001).

37. W. F. Morris, D. F. Doak, Quantitative conservation biology: theory and practice of population viability analysis., (Sinauer Associates, Sunderland, Massachusetts, 2002).

38. W. F. Frick, D. S. Reynolds, T. H. Kunz, Influence of climate and reproductive timing on demography of little brown myotis Myotis lucifugus. J Anim Ecol 79, 128–136 (2010).

39. J. D. Reichard, T. H. Kunz, White-Nose Syndrome Inflicts Lasting Injuries to the Wings of Little Brown Myotis (Myotis lucifugus). Acta Chiropterologica 11, 457–464 (2009).

40. N. B. J. S. A. Simmons, BIOLOGY-Taking Wing-Fossil and genetic findings elucidate the evolution of bats--And settle a long-standing debate over the origins of flight and echolocation. 96 (2008).

41. N. O. Therkildsen, S. R. Palumbi, Practical low-coverage genomewide sequencing of hundreds of individually barcoded samples for population and evolutionary genomics in nonmodel species. Mol Ecol Resour, (2016).

42. M. Baym et al., Inexpensive multiplexed library preparation for megabase-sized genomes. PLoS One 10, e0128036 (2015).

43. T. S. Korneliussen, A. Albrechtsen, R. Nielsen, ANGSD: Analysis of Next Generation Sequencing Data. BMC bioinformatics 15,(2014).

44. R. Nielsen, T. Korneliussen, A. Albrechtsen, Y. Li, J. Wang, SNP calling, genotype calling, and sample allele frequency estimation from New-Generation Sequencing data. PLoS One 7, e37558 (2012).

45. C. Alex Buerkle, Z. Gompert, Population genomics based on low coverage sequencing: how low should we go? Molecular ecology 22, 3028–3035 (2013).

46. E. C. Anderson, H. J. Skaug, D. J. Barshis, Next-generation sequencing for molecular ecology: a caveat regarding pooled sampled. Molecular ecology 23, 502–512 (2014).

47. D. J. Cutler, J. D. Jensen, To pool, or not to pool? Genetics 186, 41–43 (2010).

48. B. Langmead, S. L. Salzberg, Fast gapped-read alignment with Bowtie 2. Nature methods 9, 357–359 (2012).

49. A. R. Quinlan, I. M. Hall, BEDTools: a flexible suite of utilities for comparing genomic features. Bioinformatics 26, 841–842 (2010).

50. T. Derrien et al., Fast computation and applications of genome mappability. PLoS One 7, e30377 (2012).

51. A. Smit, R. Hubley, P. Green. (2013-2015).

52. A. S. Mikheyev, M. M. Tin, J. Arora, T. D. Seeley, Museum samples reveal rapid evolution by wild honey bees exposed to a novel parasite. Nature communications 6, 7991 (2015).

53. D. J. Prince et al., The Evolutionary Basis of Premature Migration in Pacific Salmon Highlights the Utility of Genomics for Informing Conservation. Science Advances, (2017).

54. J. Meisner, A. Albrechtsen, (2018).

55. R. A. Fisher, On the Dominance Ratio. Proceedings of the Royal Society Edinburgh 42, 321–431 (1922).

56. S. Wright, Evolution in Mendelian Populatins. Genetics 2, 97–159 (1930).

57. R. S. Waples, G. Luikart, J. R. Faulkner, D. A. Tallmon, Simple life-history traits explain key effective population size ratios across diverse taxa. Proceedings. Biological sciences /The Royal Society 280, 20131339 (2013).

58. B. V. North, D. Curtis, P. C. Sham, A note on the calculation of empirical P values from Monte Carlo procedures. Am J Hum Genet 71, 439–441 (2002).

59. M. Dewey, metap: meta-analysis of significance values. R package version 1.1., (2019).

60. K. G. Welinder et al., Biochemical Foundations of Health and Energy Conservation in Hibernating Free-ranging Subadult Brown Bear Ursus arctos. J Biol Chem 291, 22509–22523 (2016).

61. J. A. Badway, J. D. Baleja, Reps2: A cellular signaling and molecular trafficking nexus. The International Journal of Biochemistry & Cell Biology 43, 1660–1663 (2011).

62. N. Hoshina et al., Protocadherin 17 Regulates Presynaptic Assembly in Topographic Corticobasal Ganglia Circuits. Neuron 78, 839–854 (2013).

63. S. Hirano, S. Suzuki, C. Redies, The Cadherin Superfamily in Neural Development: Diversity, Funciton and Interaction with Other Molecules. Frontiers in Bioscience 8, 306–356 (2003).

64. H. Chang et al., The protocadherin 17 gene affects cognition, personality, amygdala structure and function, synapse development and risk of major mood disorders. Mol Psychiatry 23, 400–412 (2018).

65. A. S. Axelsson et al., Sox5 regulates beta-cell phenotype and is reduced in type 2 diabetes. Nature communications 8, 15652 (2017).

66. M. C. Kruer et al., Defective FA2H leads to a novel form of neurodegeneration with brain iron accumulation (NBIA). Annals of Neurology 68, 611–618 (2010).

67. S. A. Schneider, K. P. Bhatia, Three faces of the same gene: FA2H links neurodegeneration with brain iron accumulation, leukodystrophies, and hereditary spastic paraplegias. Annals of Neurology 68,(2010).

68. E. Stergiakouli et al., Genome-wide association study of height-adjusted BMI in childhood identifies functional variant in ADCY3. Obesity (Silver Spring) 22, 2252–2259 (2014).

69. S. Saeed et al., Loss-of-function mutations in ADCY3 cause monogenic severe obesity. Nature genetics 50, 175–179 (2018).

70. N. Grarup et al., Loss-of-function variants in ADCY3 increase risk of obesity and type 2 diabetes. Nature genetics 50, 172–174 (2018).

71. X. Zhang, J. Zara, R. K. Siu, K. Ting, C. Soo, The Role of NELL-1, a Growth Factor Associated with Craniosynostosis, in Promoting Bone Regeneration. Journal of Dental Research 89, 865–878 (2010).

72. S. B. Doty, E. A. Nunez, Activation of Osteoclasts and the Repopulation of Bone Surfaces Following Hibernation in the Bat, Myotis lucifugus. The Anatomical Record 213, 481–495 (1985).

73. G. B. Christiansen et al., The sorting receptor SorCS3 is a stronger regulator of glutamate receptor functions compared to GABAergic mechanisms in the hippocampus. Hippocampus 27, 235–248 (2017).

74. T. Breiderhoff et al., Sortilin-related receptor SORCS3 is a postsynaptic modulator of synaptic depression and fear extinction. PLoS One 8, e75006 (2013).

75. L. Qu et al., Association between polymorphisms in RAPGEF1, TP53, NRF1 and type 2 diabetes in Chinese Han population. Diabetes Res Clin Pract 91, 171–176 (2011).

76. V. Radha, A. Rajanna, R. K. Gupta, K. Dayma, T. Raman, The guanine nucleotide exchange factor, C3G regulates differentiation and survival of human neuroblastoma cells. Journal of Neurochemistry 107, 1424–1435 (2008).

77. J. M. Keaton et al., Genome-wide interaction with the insulin secretion locus MTNR1B reveals CMIP as a novel type 2 diabetes susceptibility gene in African Americans. Genet Epidemiol 42, 559–570 (2018).

78. D. Wu, A. K. Murashov, MicroRNA-431 regulates axon regeneration in mature sensory neurons by targeting the Wnt antagonist Kremen1. Front Mol Neurosci 6, 35 (2013).

79. S. P. Ross et al., miRNA-431 Prevents Amyloid-beta-Induced Synapse Loss in Neuronal Cell Culture Model of Alzheimer’s Disease by Silencing Kremen1. Front Cell Neurosci 12, 87 (2018).

