## Supplemental figures and tables for "Genomic signatures of evolutionary rescue in bats surviving white-nose syndrome"

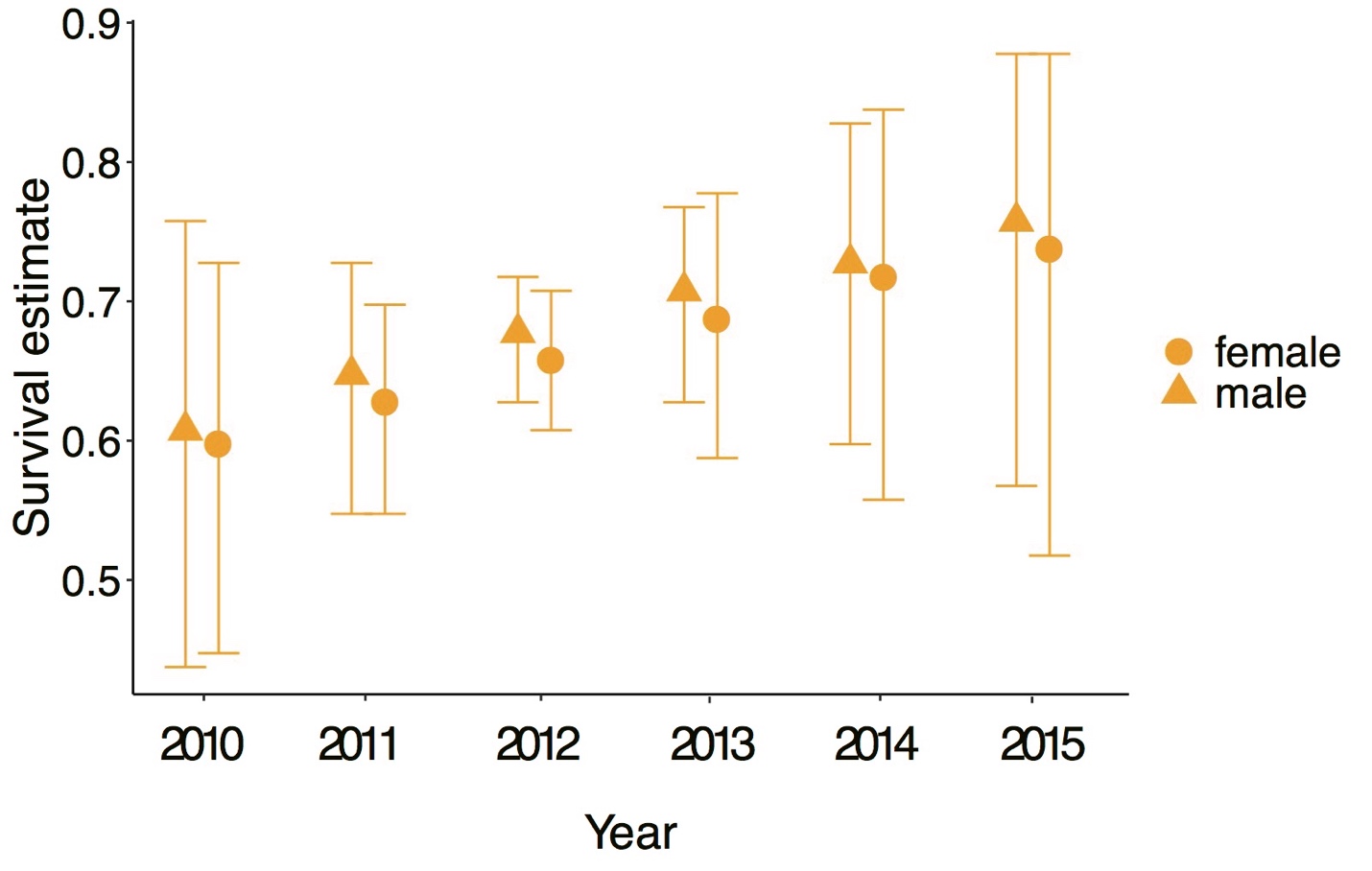

**Figure S1.** **Survival increased for both male and female little brown bat at Hibernia mine across the study period.** Survival rates in 2015 were equivalent to the published pre-WNS average of 0.76 (*38*). Error bars represent 95% confidence intervals.

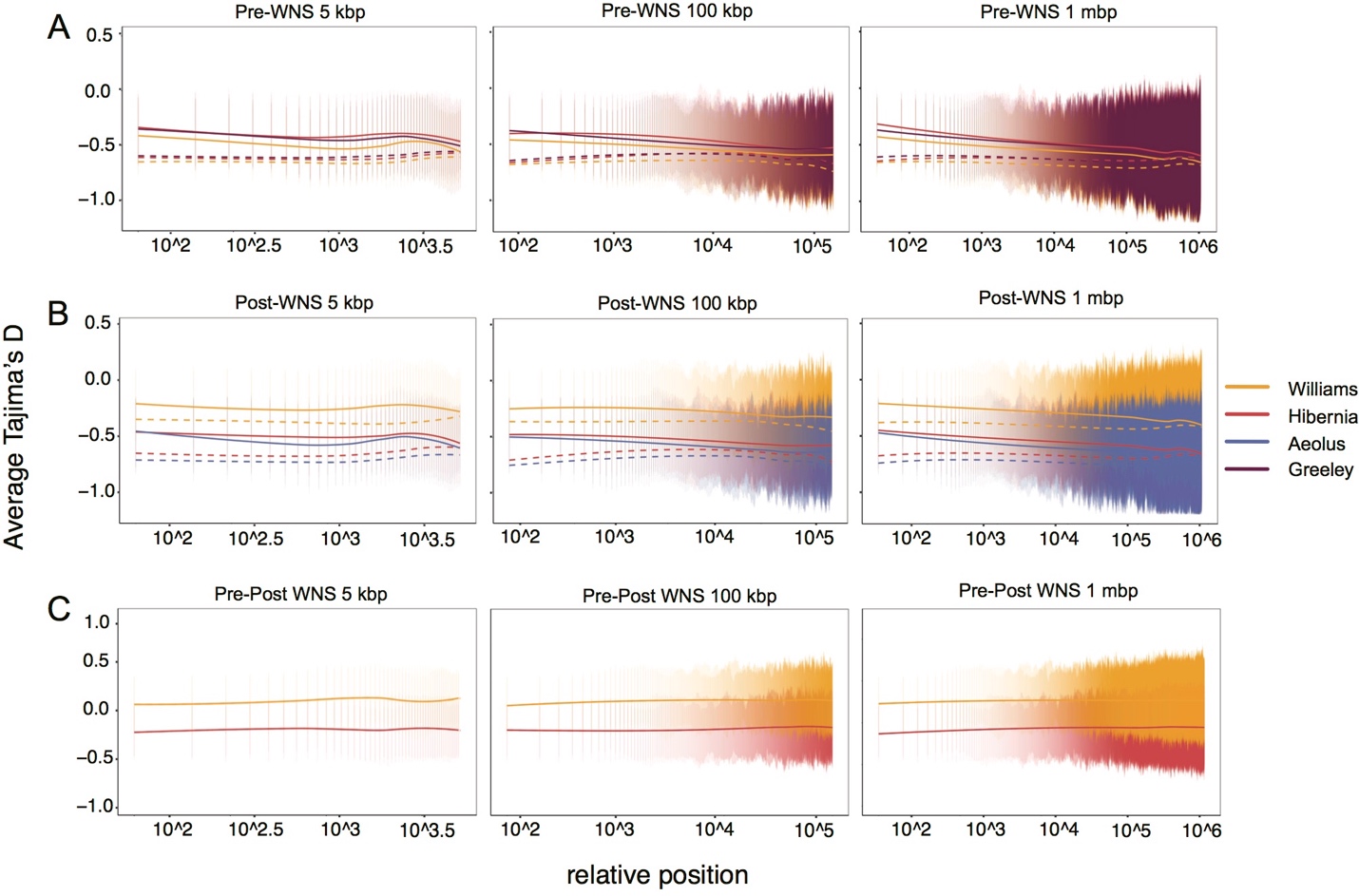

Fig. S2. Regions surrounding candidate SNPs under selection do not show larger changes in Tajima’s D between pre- and post-WNS populations than regions far from candidate SNPs. Tajima’s D was calculated in 100 basepair sliding windows surrounding candidate SNPs (solid lines) and randomly chosen non-candidate SNPs (dotted lines) in A) pre-WNS populations, B) post-WNS populations, and C) the difference between pre and post-WNS for Hibernia and Williams populations. X-axis is the distance in basepairs from SNPs of interest. Panels display the same results at different genome scales, from left to right: 5 kilobasepairs, 100 kilobasepairs, 1 megabasepairs. Error bars are standard deviations.

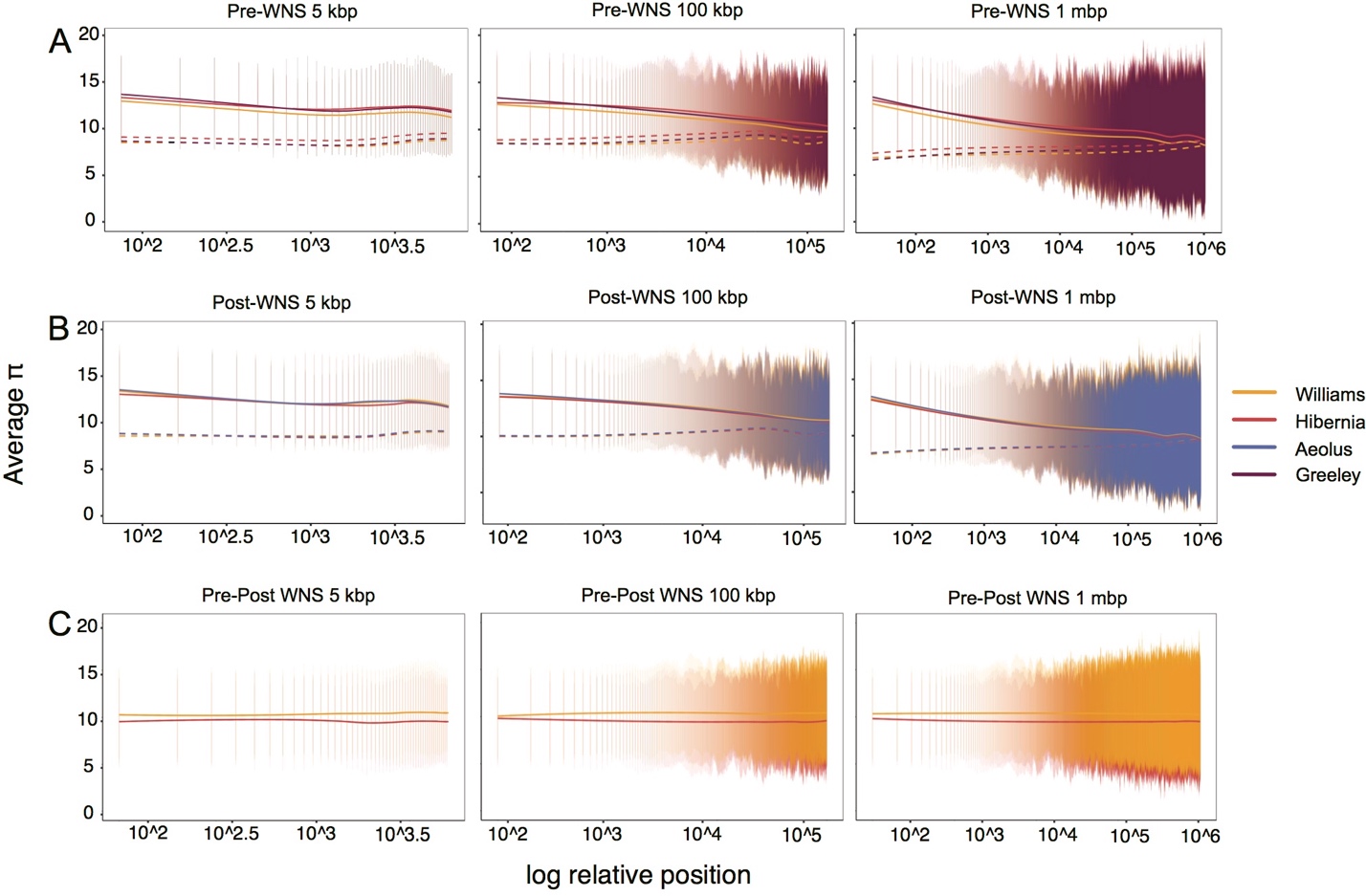

Fig. S3. Regions surrounding candidate SNPs under selection do not show changes in π between pre- and post-WNS populations. π was calculated in 100 basepair sliding windows surrounding candidate SNPs (solid lines) and randomly chosen non-candidate SNPs (dotted lines) in A) pre-WNS populations, B) post-WNS populations, and C) the difference between pre and post-WNS for Hibernia and Williams populations. X-axis is the distance in basepairs from SNPs of interest. Panels display the same results at different genome scales, from left to right: 5 kilobasepairs, 100 kilobasepairs, 1 megabasepairs. Error bars are standard deviations.

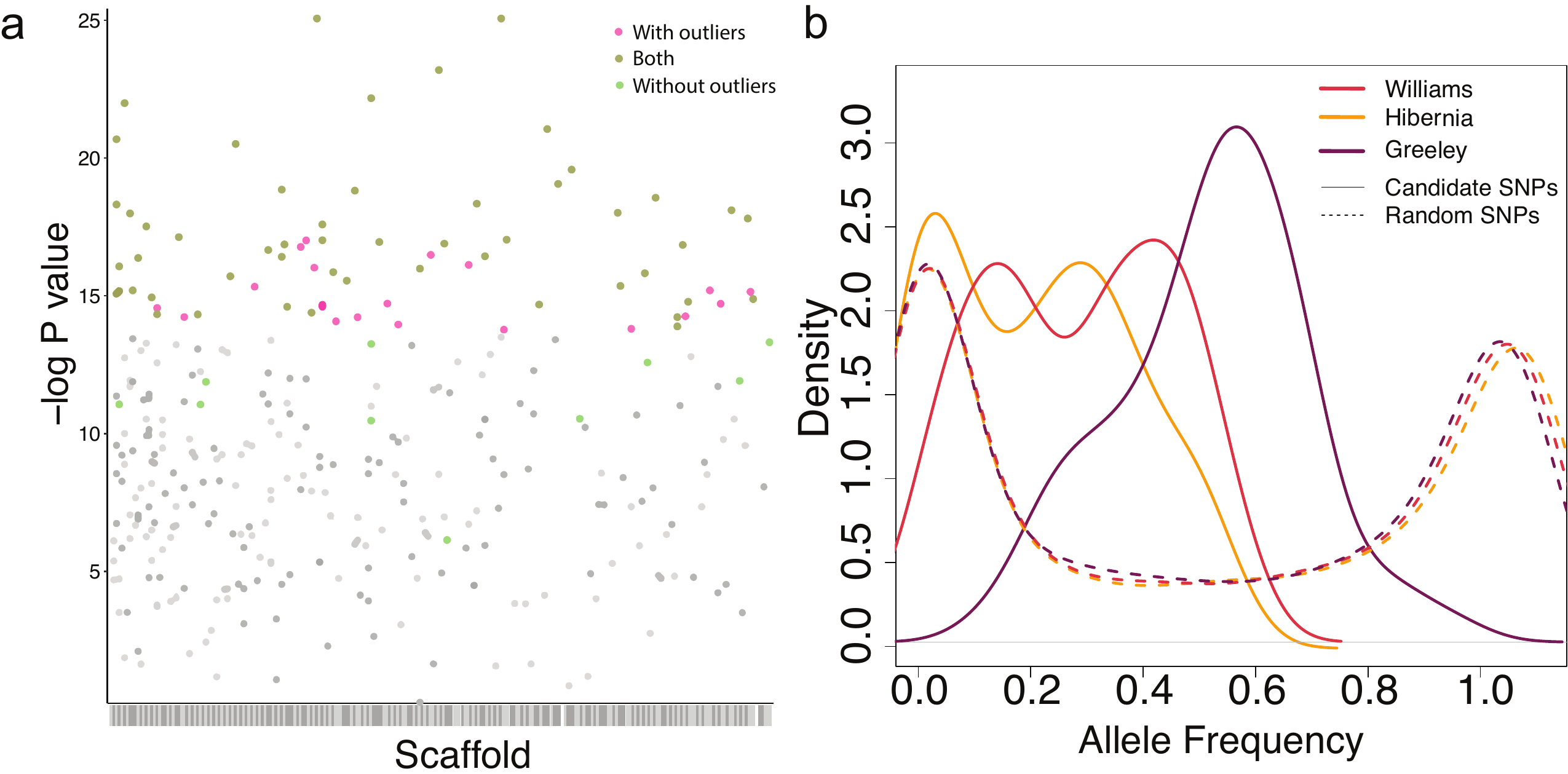

**Fig. S4.** **The addition of two pre-WNS outlier bats from Williams and two from Greeley (Fig. 2) only slightly altered the significant SNPs and pre-WNS allele frequencies.** a) The number of significant SNPs increased, but the majority remained significant both with and without outliers. The figure shows –log10 P values for SNPs with a positive change in allele frequency greater than 0.5 in both Hibernia and Williams. Significant SNPs (FDR=0.2) as determined with (99 SNPs), without (85 SNPs) and both with and without (70 SNPs) the two outlier bats are highlighted. b) Density plots of allele frequencies for significant candidate SNPs (solid lines) and random SNPs (dotted lines) at each site pre-WNS with outlier bats included.

**Table S1. Average depth of coverage per sample across the entire genome ranged from 0.61 to 8.14.**

| **Sample Name** | **Site** | **Timepoint** | **Average Depth** |
| --- | --- | --- | --- |
| NJ_E_01 | Hibernia | Post-WNS | 0.63 |
| NJ_E_02 | Hibernia | Post-WNS | 1.97 |
| NJ_E_04 | Hibernia | Post-WNS | 1.44 |
| NJ_E_05 | Hibernia | Post-WNS | 3.19 |
| NJ_E_06 | Hibernia | Post-WNS | 6.54 |
| NJ_E_07 | Hibernia | Post-WNS | 8.14 |
| NJ_E_08 | Hibernia | Post-WNS | 1.2 |
| NJ_E_10 | Hibernia | Post-WNS | 2.81 |
| NJ_E_13 | Hibernia | Post-WNS | 3.08 |
| NJ_E_14 | Hibernia | Post-WNS | 5.43 |
| NJ_E_15 | Hibernia | Post-WNS | 3.1 |
| NJ_E_16 | Hibernia | Post-WNS | 3.45 |
| NJ_E_18 | Hibernia | Post-WNS | 3.87 |
| NJ_E_21 | Hibernia | Post-WNS | 4.11 |
| NJ_E_23 | Hibernia | Post-WNS | 2.84 |
| NJ_E_24 | Hibernia | Post-WNS | 2.65 |
| NJ_E_25 | Hibernia | Post-WNS | 1.62 |
| NJ_E_26 | Hibernia | Post-WNS | 3.41 |
| NJ_E_28 | Hibernia | Post-WNS | 1.33 |
| NJ_E_29 | Hibernia | Post-WNS | 1.17 |
| NJ_E_30 | Hibernia | Post-WNS | 2.47 |
| NJ_U_01 | Hibernia | Pre-WNS | 1.81 |
| NJ_U_02 | Hibernia | Pre-WNS | 3.06 |
| NJ_U_03 | Hibernia | Pre-WNS | 3.47 |
| NJ_U_05 | Hibernia | Pre-WNS | 2.1 |
| NJ_U_06 | Hibernia | Pre-WNS | 1.72 |
| NJ_U_07 | Hibernia | Pre-WNS | 0.61 |
| NJ_U_08 | Hibernia | Pre-WNS | 2.07 |
| NJ_U_09 | Hibernia | Pre-WNS | 1.71 |
| NJ_U_10 | Hibernia | Pre-WNS | 1.94 |
| NJ_U_11 | Hibernia | Pre-WNS | 1.28 |
| NJ_U_12 | Hibernia | Pre-WNS | 2.35 |
| NJ_U_15 | Hibernia | Pre-WNS | 2.16 |
| NJ_U_16 | Hibernia | Pre-WNS | 1.11 |
| NJ_U_17 | Hibernia | Pre-WNS | 1.37 |
| NJ_U_18 | Hibernia | Pre-WNS | 6.27 |
| NJ_U_19 | Hibernia | Pre-WNS | 3.3 |
| NJ_U_21 | Hibernia | Pre-WNS | 0.97 |
| NJ_U_22 | Hibernia | Pre-WNS | 0.97 |
| NJ_U_23 | Hibernia | Pre-WNS | 0.94 |
| NJ_U_25 | Hibernia | Pre-WNS | 0.99 |
| NY_E_03 | Williams | Post-WNS | 3.8 |
| NY_E_04 | Williams | Post-WNS | 1.31 |
| NY_E_05 | Williams | Post-WNS | 1.83 |
| NY_E_06 | Williams | Post-WNS | 2.76 |
| NY_E_07 | Williams | Post-WNS | 2.93 |
| NY_E_08 | Williams | Post-WNS | 1.73 |
| NY_E_09 | Williams | Post-WNS | 1.57 |
| NY_E_10 | Williams | Post-WNS | 2.44 |
| NY_E_15 | Williams | Post-WNS | 1.66 |
| NY_E_16 | Williams | Post-WNS | 2.38 |
| NY_E_17 | Williams | Post-WNS | 2.35 |
| NY_E_18 | Williams | Post-WNS | 2.24 |
| NY_E_19 | Williams | Post-WNS | 1.02 |
| NY_E_21 | Williams | Post-WNS | 1.25 |
| NY_E_22 | Williams | Post-WNS | 1.92 |
| NY_E_23 | Williams | Post-WNS | 0.98 |
| NY_E_25 | Williams | Post-WNS | 3.13 |
| NY_E_27 | Williams | Post-WNS | 1.65 |
| NY_E_28 | Williams | Post-WNS | 2.44 |
| NY_E_29 | Williams | Post-WNS | 2.15 |
| NY_E_30 | Williams | Post-WNS | 2.04 |
| NY_U_01 | Williams | Pre-WNS | 3.39 |
| NY_U_02 | Williams | Pre-WNS | 5.71 |
| NY_U_03 | Williams | Pre-WNS | 2.38 |
| NY_U_04 | Williams | Pre-WNS | 3.35 |
| NY_U_05 | Williams | Pre-WNS | 2.47 |
| NY_U_06 | Williams | Pre-WNS | 1.19 |
| NY_U_07 | Williams | Pre-WNS | 1.82 |
| NY_U_10 | Williams | Pre-WNS | 4.69 |
| NY_U_11 | Williams | Pre-WNS | 2.72 |
| NY_U_12 | Williams | Pre-WNS | 4.25 |
| NY_U_14 | Williams | Pre-WNS | 1.45 |
| NY_U_15 | Williams | Pre-WNS | 2.75 |
| NY_U_16 | Williams | Pre-WNS | 0.93 |
| NY_U_18 | Williams | Pre-WNS | 3.65 |
| NY_U_19 | Williams | Pre-WNS | 1.71 |
| NY_U_21 | Williams | Pre-WNS | 1.57 |
| NY_U_23 | Williams | Pre-WNS | 1.8 |
| NY_U_25 | Williams | Pre-WNS | 2.56 |
| NY_U_26 | Williams | Pre-WNS | 2.42 |
| NY_U_27 | Williams | Pre-WNS | 1.79 |
| NY_U_28 | Williams | Pre-WNS | 0.96 |
| NY_U_30 | Williams | Pre-WNS | 1.39 |
| VT_E_01 | Aeolus | Post-WNS | 1.48 |
| VT_E_02 | Aeolus | Post-WNS | 2.98 |
| VT_E_03 | Aeolus | Post-WNS | 1.74 |
| VT_E_04 | Aeolus | Post-WNS | 1.56 |
| VT_E_05 | Aeolus | Post-WNS | 1.77 |
| VT_E_06 | Aeolus | Post-WNS | 3.42 |
| VT_E_07 | Aeolus | Post-WNS | 2.31 |
| VT_E_08 | Aeolus | Post-WNS | 2.85 |
| VT_E_09 | Aeolus | Post-WNS | 2.1 |
| VT_E_10 | Aeolus | Post-WNS | 2.08 |
| VT_E_11 | Aeolus | Post-WNS | 2.81 |
| VT_E_12 | Aeolus | Post-WNS | 3.03 |
| VT_E_13 | Aeolus | Post-WNS | 1.89 |
| VT_E_14 | Aeolus | Post-WNS | 0.84 |
| VT_E_15 | Aeolus | Post-WNS | 2.12 |
| VT_E_16 | Aeolus | Post-WNS | 1.4 |
| VT_E_17 | Aeolus | Post-WNS | 1.95 |
| VT_E_18 | Aeolus | Post-WNS | 2.22 |
| VT_E_19 | Aeolus | Post-WNS | 2.19 |
| VT_E_21 | Aeolus | Post-WNS | 0.8 |
| VT_E_22 | Aeolus | Post-WNS | 3.34 |
| VT_E_24 | Aeolus | Post-WNS | 1.23 |
| VT_E_25 | Aeolus | Post-WNS | 1.35 |
| VT_E_26 | Aeolus | Post-WNS | 1.72 |
| VT_E_28 | Aeolus | Post-WNS | 1.94 |
| VT_E_29 | Aeolus | Post-WNS | 1.78 |
| VT_E_30 | Aeolus | Post-WNS | 1.91 |
| VT_U_01 | Greeley | Pre-WNS | 1.46 |
| VT_U_02 | Greeley | Pre-WNS | 2.44 |
| VT_U_09 | Greeley | Pre-WNS | 2.91 |
| VT_U_10 | Greeley | Pre-WNS | 3.4 |
| VT_U_11 | Greeley | Pre-WNS | 5.71 |
| VT_U_12 | Greeley | Pre-WNS | 4.94 |
| VT_U_14 | Greeley | Pre-WNS | 3.71 |
| VT_U_15 | Greeley | Pre-WNS | 2.78 |
| VT_U_17 | Greeley | Pre-WNS | 3.08 |
| VT_U_19 | Greeley | Pre-WNS | 2.79 |
| VT_U_20 | Greeley | Pre-WNS | 1.96 |
| VT_U_22 | Greeley | Pre-WNS | 1.33 |
| VT_U_25 | Greeley | Pre-WNS | 2.65 |
| VT_U_26 | Greeley | Pre-WNS | 1.76 |
| VT_U_27 | Greeley | Pre-WNS | 2.92 |
| VT_U_29 | Greeley | Pre-WNS | 3.68 |
| VT_U_31 | Greeley | Pre-WNS | 2.79 |
| VT_U_33 | Greeley | Pre-WNS | 3.27 |
| VT_U_37 | Greeley | Pre-WNS | 2.62 |
| VT_U_40 | Greeley | Pre-WNS | 1.08 |

**Table S2.** **Most bats tagged at Hibernia were recaptured at Hibernia**. A small number of Hibernia-banded bats were found at nearby hibernacula. Three bats were found at a distant hibernaculum.

| **Location** | **Number of bands recaptured (>= 1 time)** | **Distance from first band (km)** |
| --- | --- | --- |
| Hibernia Mine | 913 | 0 |
| Mt Hope Mine | 11 | 3.81 |
| Pompton River | 1 | 17.83 |
| Williams | 1 confirmed, 2 probable | 116.89 |

**Table S3. Bats sampled at Hibernia were frequently recaptured at Hibernia.** Out of 21 bats sampled at Hibernia, 20 were banded and re-sighted multiple times during the study period. Not all bands can be identified during every survey as bats frequently hide in drill holes within the mine.

| **Sample name** | **Band #** | **Year banded** | **# years recaptured at Hibernia (2010-2017)** |
| --- | --- | --- | --- |
| NJ_E_01 | A01225 | 2011 | 6 |
| NJ_E_02 | A01108 | 2011 | 5 |
| NJ_E_04 | A00573 | 2010 | 7 |
| NJ_E_05 | A00286 | 2010 | 3 |
| NJ_E_06 | A01053 | 2011 | 2 |
| NJ_E_07 | A00549 | 2010 | 3 |
| NJ_E_08 | A00277 | 2010 | 6 |
| NJ_E_10 | A02152 | 2011 | 2 |
| NJ_E_13 | A0347 | 2012 | 2 |
| NJ_E_14 | A03547 | 2012 | 3 |
| NJ_E_15 | NA |  |  |
| NJ_E_16 | A05586 | 2015 | 1 |
| NJ_E_18 | A03468 | 2012 | 4 |
| NJ_E_21 | A01546 | 2011 | 4 |
| NJ_E_23 | A01647 | 2011 | 5 |
| NJ_E_24 | A01145 | 2011 | 6 |
| NJ_E_25 | A01139 | 2011 | 4 |
| NJ_E_26 | A03520 | 2012 | 4 |
| NJ_E_28 | A00591 | 2010 | 4 |
| NJ_E_29 | A01303 | 2011 | 4 |
| NJ_E_30 | A05162 | 2014 | 1 |

**Table S4. At least 50% of individuals would need to immigrate to achieve the observed post-WNS allele frequencies in selected SNPs.** In order to determine the independence of bats at Williams and Hibernia hibernacula, we calculated the percent required to immigrate from one to the other in order to create the change in allele frequencies observed for our candidate SNPs (denoted by position on scaffold). NA indicates that no amount of immigration from the other hibernaculum would achieve observed allele frequencies.

| **Scaffold** | **Position** | **Williams to Hibernia** | **Hibernia to Williams** |
| --- | --- | --- | --- |
| GL429767 | 16839644 | NA | 80% |
| GL429767 | 22649683 | 61% | NA |
| GL429767 | 38456248 | NA | 75% |
| GL429767 | 46534125 | 81% | NA |
| GL429767 | 51542832 | 98% | NA |
| GL429768 | 29962249 | NA | 96% |
| GL429768 | 34577793 | 83% | NA |
| GL429768 | 3921142 | 77% | NA |
| GL429770 | 8292792 | 70% | NA |
| GL429770 | 8507307 | 93% | NA |
| GL429772 | 14993622 | 94% | NA |
| GL429772 | 8716095 | 96% | NA |
| GL429773 | 7204887 | 90% | NA |
| GL429775 | 5556309 | 88% | NA |
| GL429778 | 3561000 | 94% | NA |
| GL429781 | 7594948 | NA | 82% |
| GL429783 | 14617697 | 89% | NA |
| GL429783 | 665407 | 99% | NA |
| GL429784 | 1229639 | 89% | NA |
| GL429785 | 3216149 | 85% | NA |
| GL429792 | 753145 | 86% | NA |
| GL429794 | 5707870 | 93% | NA |
| GL429795 | 3907289 | 70% | NA |
| GL429801 | 6986210 | NA | 75% |
| GL429809 | 7390034 | NA | 99% |
| GL429816 | 3229647 | NA | 98% |
| GL429819 | 365354 | 69% | NA |
| GL429828 | 1501267 | 90% | NA |
| GL429828 | 3004686 | 96% | NA |
| GL429835 | 2966433 | 78% | NA |
| GL429835 | 2966452 | NA | 97% |
| GL429841 | 3522450 | NA | 79% |
| GL429841 | 612532 | 66% | NA |
| GL429842 | 4124241 | 74% | NA |
| GL429845 | 2091046 | 75% | NA |
| GL429848 | 2603829 | NA | 93% |
| GL429850 | 1183154 | 72% | NA |
| GL429852 | 463102 | NA | 91% |
| GL429855 | 4401152 | 90% | NA |
| GL429857 | 134612 | 77% | NA |
| GL429859 | 300331 | 89% | NA |
| GL429861 | 293372 | 69% | NA |
| GL429861 | 3065399 | 91% | NA |
| GL429861 | 4317517 | NA | 98% |
| GL429861 | 4317523 | 99% | NA |
| GL429867 | 2575249 | 85% | NA |
| GL429873 | 2056629 | 67% | NA |
| GL429879 | 384336 | 92% | NA |
| GL429880 | 42824 | 62% | NA |
| GL429885 | 2043530 | 87% | NA |
| GL429888 | 3508435 | NA | 70% |
| GL429891 | 266596 | 87% | NA |
| GL429893 | 1767654 | 96% | NA |
| GL429906 | 1866953 | NA | 77% |
| GL429910 | 2414078 | NA | 96% |
| GL429920 | 2255691 | NA | 85% |
| GL429927 | 1241213 | 84% | NA |
| GL429929 | 2943933 | 86% | NA |
| GL429938 | 2500270 | 92% | NA |
| GL429946 | 1039079 | NA | 78% |
| GL429947 | 1200485 | 51% | NA |
| GL429950 | 2444879 | 88% | NA |
| GL429955 | 1484586 | 73% | NA |
| GL429962 | 689293 | 74% | NA |
| GL429965 | 1956978 | 94% | NA |
| GL429968 | 2123897 | 67% | NA |
| GL429969 | 109273 | 64% | NA |
| GL429982 | 1303809 | 88% | NA |
| GL429998 | 99324 | 95% | NA |
| GL430008 | 1396110 | NA | 69% |
| GL430018 | 180278 | 97% | NA |
| GL430029 | 1661413 | 79% | NA |
| GL430101 | 496908 | 85% | NA |
| GL430121 | 936584 | 85% | NA |
| GL430128 | 735347 | 93% | NA |
| GL430132 | 776142 | 80% | NA |
| GL430156 | 916661 | 94% | NA |
| GL430166 | 11932 | 85% | NA |
| GL430239 | 60325 | 77% | NA |
| GL430271 | 326498 | NA | 92% |
| GL430271 | 525210 | 96% | NA |
| GL430279 | 607165 | 72% | NA |
| GL430281 | 514067 | NA | 96% |
| GL430338 | 350547 | NA | 83% |
| GL430384 | 252240 | 85% | NA |
| GL430420 | 180654 | 67% | NA |
| GL430482 | 173352 | NA | 96% |
| GL430686 | 91034 | 87% | NA |
| GL430696 | 55439 | 81% | NA |
| GL430746 | 9663 | 89% | NA |
| GL430834 | 44020 | 97% | NA |
| GL430870 | 25568 | 64% | NA |

**Table S5. Sixty-two significant SNPs were identified on multiple scaffolds throughout the genome.** Significance was determined using the Benjamini-Hochberg critical value calculated from p-values combined using Fisher’s sum log method. Gene annotations are listed when known.

| Ensembl Scaffold | NCBI Scaffold | Location on scaffold | Major allele | Minor allele | Combined p-value | Benjamini-Hochberg critical value | Annotation |
| --- | --- | --- | --- | --- | --- | --- | --- |
| GL429767 | NW_005871048.1 | 16839644 | T | A | 7.51E-07 | 8.49E-07 | FBXL17 |
| GL429767 | NW_005871048.1 | 22649683 | T | A | 6.76E-08 | 3.22E-07 |  |
| GL429767 | NW_005871048.1 | 46534125 | C | T | 1.64E-09 | 1.02E-07 |  |
| GL429768 | NW_005871049.1 | 3921142 | G | C | 5.16E-07 | 7.32E-07 |  |
| GL429768 | NW_005871049.1 | 29962249 | A | G | 1.05E-07 | 3.81E-07 |  |
| GL429768 | NW_005871049.1 | 9041952 | A | G | 8.73E-07 | 8.93E-07 |  |
| GL429770 | NW_005871051.1 | 8507307 | A | C | 2.86E-11 | 1.46E-08 |  |
| GL429772 | NW_005871053.1 | 8716095 | T | A | 1.31E-08 | 2.06E-07 |  |
| GL429772 | NW_005871053.1 | 14993622 | T | C | 4.51E-07 | 6.88E-07 |  |
| GL429773 | NW_005871054.1 | 7204887 | C | T | 8.35E-07 | 8.78E-07 |  |
| GL429775 | NW_005871056.1 | 5556309 | A | C | 1.98E-07 | 5.27E-07 |  |
| GL429778 | NW_005871059.1 | 3561000 | G | A | 6.21E-08 | 3.09E-07 |  |
| GL429781 | NW_005871062.1 | 7594948 | A | T | 3.28E-07 | 6.29E-07 | MASP1 |
| GL429783 | NW_005871064.1 | 665407 | C | T | 6.40E-07 | 7.91E-07 |  |
| GL429792 | NW_005871073.1 | 753145 | C | A | 3.30E-07 | 6.44E-07 |  |
| GL429801 | NW_005871082.1 | 6986210 | A | T | 9.00E-07 | 9.08E-07 |  |
| GL429802 | NW_005871083.1 | 4091453 | C | T | 1.68E-07 | 4.83E-07 | CMIP |
| GL429806 | NW_005871087.1 | 3311579 | T | C | 3.61E-07 | 6.73E-07 |  |
| GL429816 | NW_005871097.1 | 3229647 | G | A | 1.62E-07 | 4.54E-07 | REPS2 |
| GL429819 | NW_005871100.1 | 365354 | G | A | 2.86E-11 | 2.93E-08 |  |
| GL429835 | NW_005871116.1 | 2966433 | A | G | 6.65E-07 | 8.05E-07 |  |
| GL429841 | NW_005871122.1 | 3522450 | A | G | 1.81E-07 | 5.12E-07 | PCDH17 |
| GL429841 | NW_005871122.1 | 612532 | G | A | 2.14E-08 | 2.34E-07 |  |
| *GL429842* | *NW_005871123.1* | 4124241 | *A* | *G* | *5.56E-08* | *2.98E-07* |  |
| GL429845 | NW_005871126.1 | 2091046 | C | T | 7.04E-07 | 8.12E-07 | SOX5 |
| GL429855 | NW_005871136.1 | 4401152 | G | C | 5.46E-07 | 7.30E-07 | FA2H |
| GL429859 | NW_005871140.1 | 300331 | T | C | 2.86E-11 | 1.03E-08 |  |
| GL429861 | NW_005871142.1 | 293372 | A | G | 4.82E-08 | 2.88E-07 |  |
| GL429861 | NW_005871142.1 | 3065399 | A | G | 1.46E-07 | 4.63E-07 | ADCY3 |
| GL429867 | NW_005871148.1 | 2575249 | A | G | 4.62E-07 | 6.99E-07 | CEP112 |
| GL429873 | NW_005871154.1 | 2056629 | T | A | 6.79E-07 | 7.92E-07 |  |
| GL429879 | NW_005871160.1 | 384336 | G | A | 1.66E-07 | 4.94E-07 |  |
| GL429885 | NW_005871166.1 | 3445677 | T | A | 4.97E-07 | 7.10E-07 |  |
| GL429885 | NW_005871166.1 | 617822 | C | T | 3.13E-07 | 6.17E-07 |  |
| GL429885 | NW_005871166.1 | 2043530 | T | C | 1.35E-10 | 7.20E-08 |  |
| GL429888 | NW_005871169.1 | 3508435 | A | T | 1.31E-07 | 4.42E-07 |  |
| GL429910 | NW_005871191.1 | 2414078 | A | C | 2.10E-07 | 5.45E-07 | NELL1 |
| GL429927 | NW_005871208.1 | 1241213 | A | G | 9.77E-09 | 1.75E-07 |  |
| GL429929 | NW_005871210.1 | 2943933 | G | T | 1.21E-07 | 4.22E-07 | SORCS3 |
| GL429930 | NW_005871211.1 | 2586279 | C | T | 5.92E-07 | 7.40E-07 |  |
| GL429950 | NW_005871231.1 | 2444879 | G | A | 6.20E-07 | 7.61E-07 |  |
| GL429955 | NW_005871236.1 | 1484586 | G | A | 8.24E-08 | 3.60E-07 |  |
| GL429962 | NW_005871243.1 | 689293 | A | C | 2.86E-11 | 3.09E-08 |  |
| GL429965 | NW_005871246.1 | 1956978 | G | A | 4.74E-08 | 2.67E-07 | CBFA2T3 |
| GL429998 | NW_005871279.1 | 99324 | C | G | 2.16E-07 | 5.55E-07 |  |
| GL430008 | NW_005871289.1 | 1396110 | C | T | 1.09E-10 | 6.17E-08 |  |
| GL430018 | NW_005871299.1 | 180278 | T | A | 7.32E-09 | 1.65E-07 |  |
| GL430029 | NW_005871310.1 | 1661413 | A | C | 4.53E-09 | 1.44E-07 | GEMIN4 |
| GL430036 | NW_005871317.1 | 626424 | G | T | 2.82E-09 | 1.23E-07 | KREMEN1 |
| GL430101 | NW_005871382.1 | 496908 | T | C | 1.92E-08 | 2.26E-07 |  |
| GL430121 | NW_005871402.1 | 936584 | T | C | 6.79E-08 | 3.39E-07 |  |
| GL430156 | NW_005871437.1 | 916661 | C | T | 1.46E-07 | 4.73E-07 |  |
| GL430158 | NW_005871439.1 | 593841 | T | G | 2.01E-09 | 1.13E-07 |  |
| GL430166 | NW_005871447.1 | 11932 | C | T | 3.28E-07 | 6.27E-07 |  |
| GL430239 | NW_005871520.1 | 60325 | G | A | 1.76E-07 | 5.14E-07 |  |
| GL430239 | NW_005871520.1 | 60326 | T | C | 2.37E-07 | 5.96E-07 |  |
| GL430271 | NW_005871552.1 | 525210 | C | T | 8.98E-08 | 3.81E-07 | ERCC4 |
| GL430281 | NW_005871562.1 | 514067 | T | C | 2.21E-07 | 5.66E-07 |  |
| GL430482 | NW_005871763.1 | 173352 | C | T | 1.48E-08 | 2.16E-07 |  |
| GL430565 | NW_005871846.1 | 94959 | A | G | 7.85E-07 | 8.43E-07 |  |
| GL430686 | NW_005871967.1 | 91034 | C | T | 4.76E-08 | 2.78E-07 |  |
| GL430746 | NW_005872028.1 | 9663 | C | T | 3.47E-07 | 6.58E-07 | RAPGEF1 |
| GL432768 | NW_005875640.1 | 965 | C | T | 1.17E-08 | 1.95E-07 |  |

**Table S6. Significant SNPs were found in genes associated with hibernation phenotypes.** Proteins coded for by genes associated with SNPs that underwent significant changes in allele frequency. Function is listed when known.

| Gene | Protein | Putative Function |
| --- | --- | --- |
| MASP1 | Mannan-binding lectin serine protease 1 | Part of the innate immune complement pathway; less abundant in hibernating brown bears (*60*) |
| REPS2 | RalBP1-associated Eps domain-containing protein 2 | Involved in insulin regulation (*61*) |
| PCDH17 | Protocadherin-17 | Regulates pre-synaptic assembly (*62*); Cadherins are involved in synaptogenesis (*63*), changes in dendritic spines, and mood disorders (*64*) |
| SOX5 | Transcription factor SOX-5 | Reduced in type 2 diabetes (*65*) |
| FA2H | Fatty acid 2-hydroxylase | Mutations lead to neurodegeneration with brain ion accumulation (*66*) and other neuro-disorders (*67*) |
| ADCY3 | Adenylate cyclase type 3 | Strongly associated with obesity and diabetes in multiple populations (*68-70*) |
| CEP112 | Centrosomal protein 112 kDa | None |
| NELL1 | Protein kinase C-binding protein NELL1 | Involved in bone regeneration (*71*), which occurs upon arousal from hibernation in little browns (*72*) |
| SORCS3 | VPS10 domain-containing receptor SorCS3 | Involved in glutamate receptor function and post-synaptic transmission (*73*); deficient mice have reduced synaptic plasticity (*74*) |
| CBFA2T3 | Protein CBFA2T3 | None |
| GEMIN4 | Gem-associated protein 4 | None |
| RAPGEF1 | Rap guanine nucleotide exchange factor 1 | Associated with diabetes (*75*); neuron differentiation and growth (*76*) |
| CMIP | C-Maf inducing protein | Associated with type 2 diabetes susceptibility (*77*) |
| Kremen1 | Kremen protein 1 | Inhibits axon regeneration (*78*) in Alzheimer’s(*79*) |
| ERCC4 | DNA repair endonuclease XPF | None |
| FBXL17 | F-box and leucine rich repeat protein 17 | None |
